# Characterization of Nanoparticles in Suspension by Simultaneous iNTA and Fluorescence Detection with Single-Molecule Sensitivity

**DOI:** 10.64898/2026.04.09.717488

**Authors:** Shuhan Jiang, Anna D. Kashkanova, Hannarae Lee, Morgan E. C. Miller, Tobias Utikal, Alexey Shkarin, Hamed Qazvini, Vahid Sandoghdar

## Abstract

Quantitative knowledge of nanoparticle properties is desirable in a large number of scientific and technological applications, but measurements with a high degree of precision usually prove to be challenging. Among a range of available methodologies, optical techniques with single particle sensitivity are especially interesting because they can reveal intrinsic heterogeneities in a fast non-invasive manner. Recently, we presented interferometric nanoparticle tracking analysis (iNTA) as a highly sensitive label-free technique that is capable of determining the size, concentration and index of refraction of different subpopulations in a suspension mixture. Here, we enhance this method with biochemical specificity through multicolor fluorescence detection at the single-molecule sensitivity limit. We benchmark the performance of the combined technique, which we name iNTA-F, by distinguishing populations of fluorescent and non-fluorescent nanoparticles of different material, size, and fluorescence intensity, with an emphasis on the characterization of lipid vesicles and biological extracellular vesicles (EVs).

## Introduction

Nanoparticles play a central role in numerous scientific and technological contexts, such as colloidal chemistry,^1^ environmental science, ^2^ and biomedical applications.^3^ As a result, there is a strong need for quantitative approaches to nanoparticle analysis. Two of the basic features of interest are often size and material although surface coatings and their molecular properties are also of great interest. One of the conventional methods for characterization of these quantities is based on electron microscopy, providing high-resolution size information at the expense of low throughput, high cost, and operation in vacuum, which in turn compromises the native conditions of a fluidic medium.^4,5^ Scanning probe microscopy also provides high spatial resolution, but it can only access the outer surface layers of nano-objects.^6^

Optical approaches allow high-throughput non-invasive measurement of nanoparticle properties in their native environments, albeit at lower spatial resolution. Here, label-free optical detection methods are particularly desirable.^7^ And among these, resonant Rayleigh scattering provides a very sensitive fluorescence-free approach, as demonstrated by interferometric scattering (iSCAT) microscopy,^8,9^ which can reach the single-molecule limit.^10–13^ We recently introduced interferometric nanoparticle tracking analysis (iNTA),^14^ which uses iSCAT detection for tracking nanoparticles that undergo Brownian motion in a suspension. iNTA follows the workflow of nanoparticle tracking analysis (NTA),^15^ and uses the measurement of the diffusion constant obtained from single-particle trajectories to deduce particle size. In addition, it also provides independent information about particle size via particle polarizability. The higher sensitivity of iSCAT in detecting nanoparticles compared to dark-field microscopy gives iNTA a superior performance, especially in differentiating various particle species in a mixture and with size distribution extending below 50 nm.^14,16^ A particularly important application concerns biological entities such as liposomes, ^17^ viruses,^18^ extracellular vesicles (EVs),^19^ and protein aggregates,^20^ which are involved in transport, signaling, and disease.

Despite their remarkable sensitivity, scattering-based methods have the intrinsic shortcoming of lacking material specificity. Therefore, it is highly advantageous to combine them with fluorescence labeling,^21,22^ which has long been established as a workhorse for achieving molecular specificity in biological studies.^23^ In this work, we present iNTA-F as a fluorescence-enabled iNTA platform that provides simultaneous measurements of nanoparticle size, refractive index, and different binding sites with single-molecule sensitivity. We apply this approach to synthetic nanoparticles and to biological EVs, illustrating its ability to track freely diffusing particles while quantitatively resolving their molecular and physical heterogeneity.

## Results

### Setup Characterization and Calibration

Figure 1a shows the schematics of a custom-built wide-field microscope that integrates iNTA with two separate fluorescence channels. In brief, we used a laser with a wavelength of 638 nm for iSCAT detection as well as two excitation lasers at wavelengths of 488 nm and 561 nm for detecting fluorescence emission in the ranges of 500–550 nm (green channel) and 575–625 nm (red channel), respectively. To avoid spectral cross talk between the two fluorescence channels, the excitation lasers were interlaced in time. We used an Arduino microcontroller as frequency divider to synchronize the fast camera frame rate (5000 Hz) in the iSCAT channel with the excitation windows of the fluorescence channels at a significantly slower frame rate (10–100 Hz), dictated by the respective cameras.

**Figure 1:**
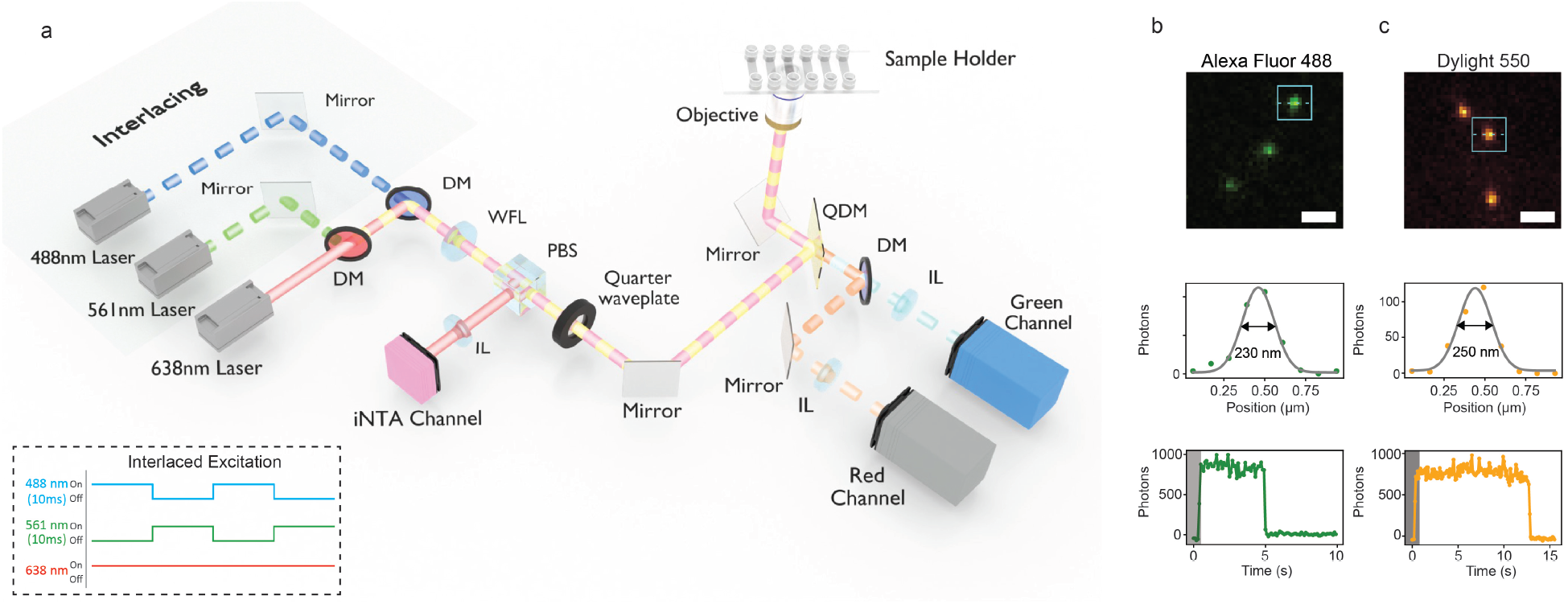
(a) Schematics of an iNTA-F setup. DM, dichroic mirror; WFL, wide-field lens; PBS, polarizing beam splitter; QDM, quad-band dichroic mirror; IL, imaging lens. The inset shows the interlaced excitation scheme, where 488 nm and 561 nm fluorescence channels are alternated (10–100 frames per second, fps), while the 638 nm laser remains continuously on for high-speed iNTA acquisition (5000 fps). (b,c) Single-molecule fluorescence characterization of Alexa Fluor 488 (b) and DyLight 550 (c) which were spin-coated on a cover glass. Top panels show representative point-spread functions (PSFs) of single molecules. Middle panels display experimental fluorescence intensity profiles with corresponding Gaussian fits. Bottom panels plot representative stepwise photobleaching traces. Scale bars, 1 µm.

To assess the fluorescence sensitivity of the setup, we imaged single Alexa Fluor 488 and DyLight 550 dye molecules on coverslips exposed to air (Figure 1b, c). Their emission spots were well described by two-dimensional (2D) Gaussian profiles with full width at half-maximum (FWHM) of 230 nm and 250 nm, respectively (top and middle panels). Stepwise photobleaching confirmed that the signals originated from individual emitters (bottom panels).

### Quantitative fluorescence characterization of single nanoparticles

To benchmark the performance of the system in an aqueous medium, we sparsely immobilized liposomes on glass coverslips. Liposomes are widely used as drug-delivery carriers in cancer therapy, vaccine formulation, and gene delivery.^24^ These are highly suited as test nanoparticles because lipid membranes can be readily functionalized with fluorescent dyes. Here, we used MemBright 488.

The number of fluorophores on a nanoparticle can be quantified if one knows the average signal from one fluorophore. To achieve this, we examined stepwise photobleaching traces from individual particles until the complete loss of signal. In the main body of Figure 2a, we present three examples for the decay of the fluorescence signal (left vertical axis) and the number of fluorophores that correspond to it (right vertical axis). The insets in Figure 2a show the iSCAT and fluorescence images and the final photobleaching steps.

**Figure 2:**
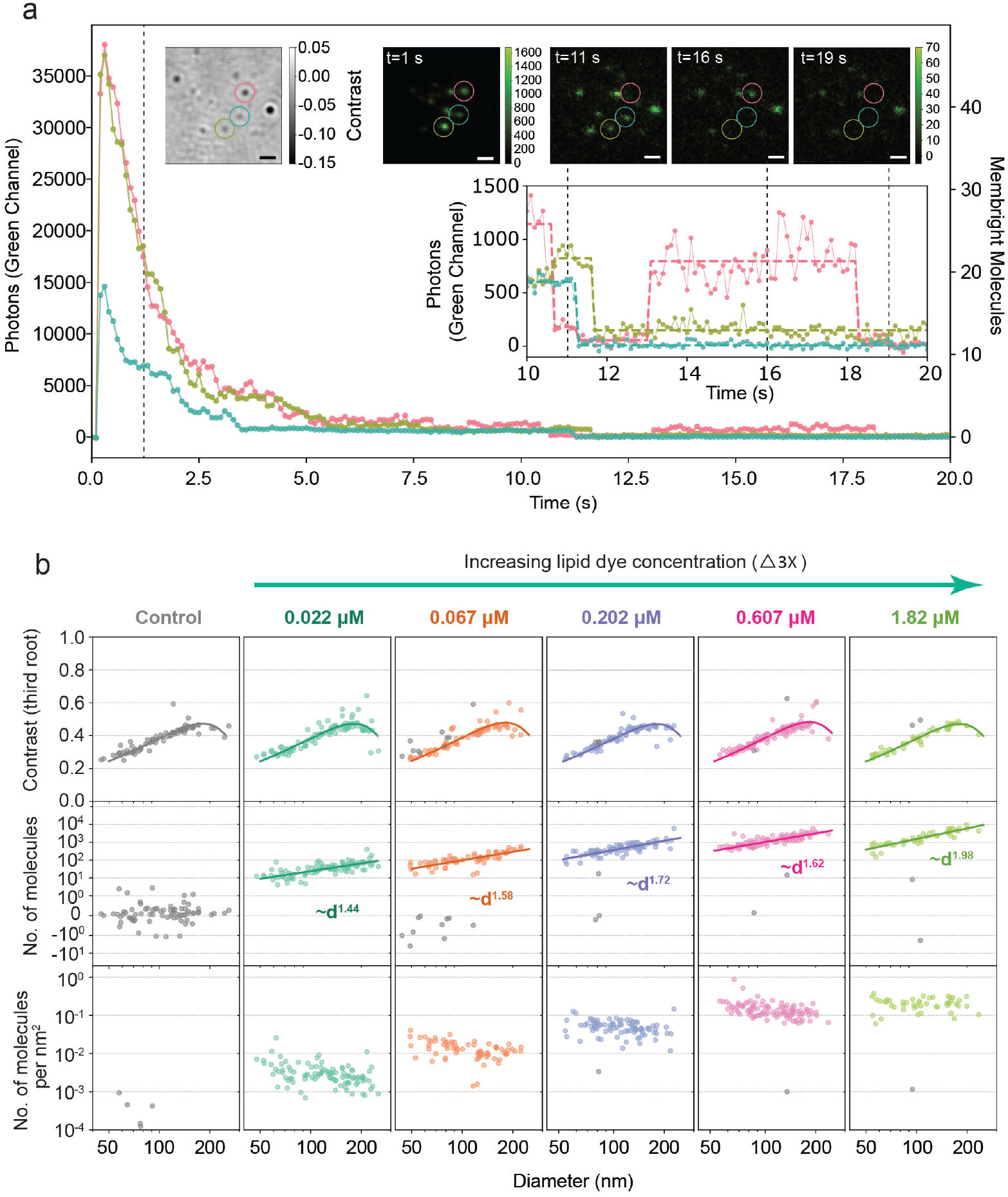
Quantitative fluorescence characterization of MemBright 488–labelled liposomes. (a) Stepwise photobleaching analysis of individual liposomes immobilized on a coverslip. Main panel shows the fluorescence intensity over time in the 488 Fluo channel. The inset displays the final 10 s where discrete photobleaching steps take place. Corresponding iSCAT and fluorescence images of a representative liposome at 1 s, 11 s, 16 s, and 19 s. Time points are indicated by dashed lines. (b) iNTA-F studies of diffusing liposomes with different dye concentrations. (Top) Conventional iNTA scatter plots of the third root of the iSCAT contrast versus particle diameter. The left column represents ‘control’ data from unlabled liposomes. Five dye labeling concentrations are indicated above each panel legend. Contours represent fits to a core–shell model with a shell refractive index (RI) of 1.45, thickness of 6 nm, and a core RI of 1.33. (Middle) Number of dye molecules per liposome as a function of particle diameter. (Bottom) Dye molecule density (number of molecules per nm^2^) as a function of liposome size, showing saturation at high labeling concentrations.

Next, we extended our measurements to freely diffusing liposomes in milli-Q water. As individual liposomes crossed the laser beam at about 2 *µ*m above the cover glass, their trajectory was imaged, allowing us to analyze their fluorescence strength. In this configuration, the collection efficiency for a randomly oriented dipole moment is reduced from 68% at the water-glass interface to about 42%. Furthermore, diffusing particles can already get excited in the tails of the laser beam profile and along the axial direction before entering our imaging field of view. This leads to faster photobleaching. Furthermore, if particles that have previously been exposed to laser light within the field of view return, they will be more prone to photobleaching since the rate for this process is correlated to the probability of visiting the excited state.^25^

To reduce the effect of photobleaching, we continuously refreshed the sample volume by introducing a gentle flow, in which particles traversed the illuminated Gaussian region with a full width at half-maximum of *∼*4 *µ*m on a time scale of seconds, comparable to or shorter than diffusion-driven re-entry. Under these conditions, particle size estimates deduced from the diffusion constant remained unchanged, while fluorescence photon counts per particle were improved.

The top left panel in Figure 2b presents a conventional iNTA scatter plot for an unlabeled sample, where the third root of the iSCAT contrast is plotted against the particle diameter determined from diffusion analysis. The signal rises linearly at first because the iSCAT contrast is proportional to the particle polarizability, which grows as *d*^3^ for small particles (*kd ≪* 1, where *k* is the wave vector and *d* is the diameter). For larger particles, one has to consider the full Mie scattering theory. This also becomes important for core-shell particles such as liposomes and extracellular vesicles, for which the scattering strength is influenced by the shell thickness and refractive index in addition to the properties of the inner core.^14,26^ The change of curvature in the solid fit curves obtained from full Mie calculations put into evidence this change of paradigm.

In the top row of Figure 2b, we also present iNTA data from five sample sets in which liposomes were labelled at different dye concentrations. The observation that the iNTA plots remain nearly identical, lets us conclude that the presence of the lipid dye did not alter the size or the iSCAT contrast of the particles. Since each particle was also tracked in the fluorescence channel, we can correlate its diameter extracted from conventional iNTA measurements with its fluorescence signal. The data in the middle panels of Figure 2b displays the number of dye molecules per particle extracted from the fluorescence signal, using the calibration obtained from the photobleaching analysis. We find that the power law exponent for dye number versus particle diameter increased from 1.44 to nearly 2 with increasing dye concentration. This suggests that dye molecules, which were added after liposome formation in our preparation procedure, are more easily admitted in smaller liposomes, possibly due to their higher membrane curvature. As the dye concentration is increased, dye insertion reaches a uniform coverage, saturating to about one molecule/nm^2^ (see bottom panel of Figure 2b). This intriguing observation merits further quantitative investigations, which go beyond the scope of our current work.

### Dual-color iNTA-F

We further tested the system with a mixed sample of 30 nm gold nanoparticles, as well as 40 nm red and 100 nm green FluoSpheres. In Figure 3a, we show snap shots of diffusing particles simultaneously imaged in the conventional iNTA and two fluorescence channels. Both the 488 nm and 561 nm excitation lasers were set to deliver a power of approximately 80 *µ*W to the sample. Under these conditions, a single green FluoSphere typically yielded *∼* 10^5^ detected photons per frame in the green channel, while the red FluoSpheres yielded *∼* 10^4^ photons per frame in the red channel. Interlaced laser excitation allows us to distinguish different fluorophores with considerably fewer false positive events than solely relying on spectral filtering of their emission. Figure 3b presents the outcome of the analysis from a total of 163 particles. The left panel distinguishes three different populations in the conventional iNTA scatter plot, and the right panel confirms that these three distinct populations are also well separated by their fluorescence intensities.

**Figure 3:**
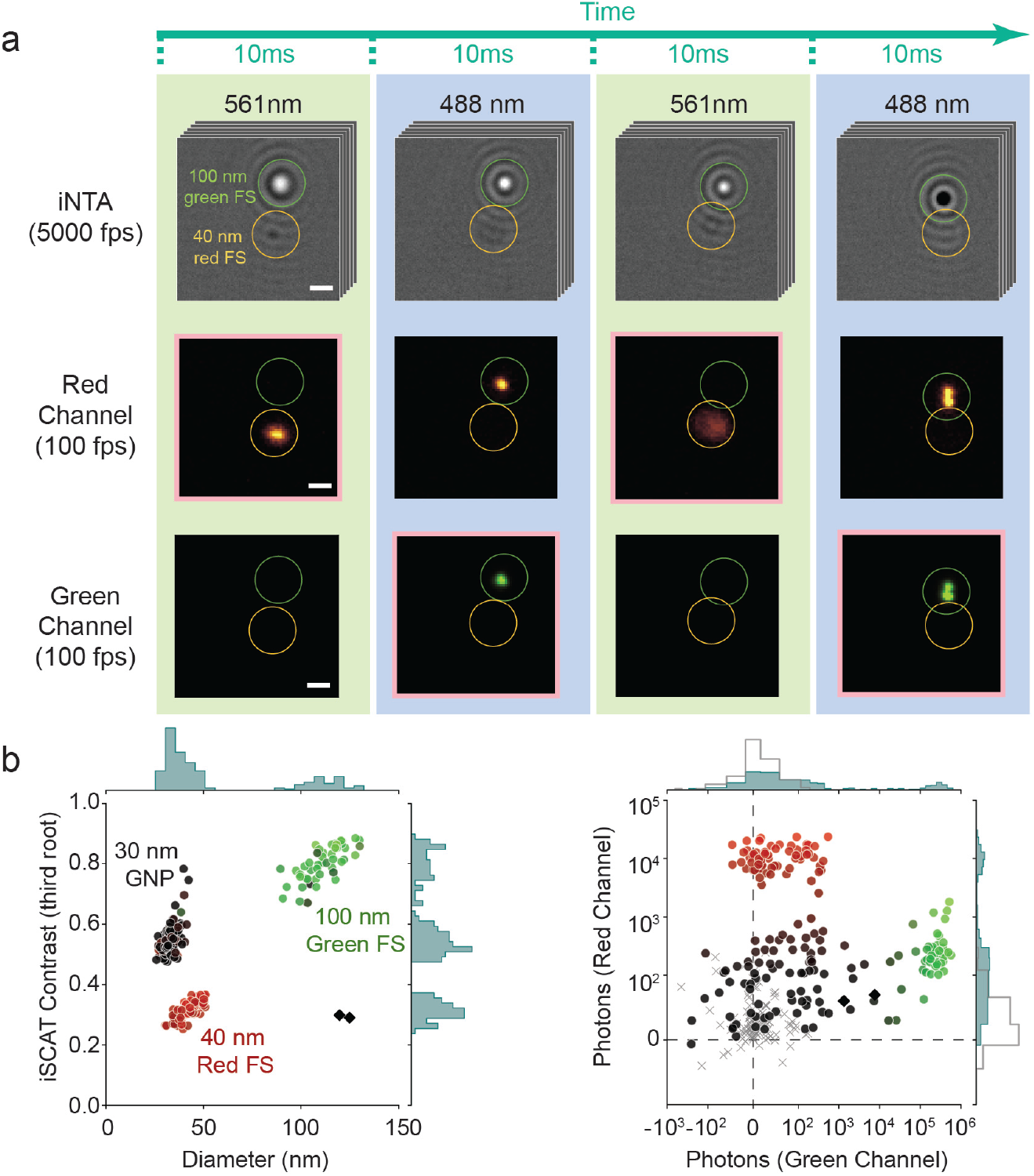
Simultaneous iNTA and fluorescence measurements of diffusing particles. (a) Simultaneous iNTA (top, 5000 fps) and fluorescence imaging (middle/bottom, 100 fps) of a mixed sample containing 100 nm green and 40 nm red FluoSpheres. Four consecutive 10 ms time intervals are displayed to illustrate the interlaced excitation of the 488 nm and 561 nm lasers. Frames used for fluorescence quantification (active excitation) are highlighted in pink. The contrast reversal in iSCAT images indicates particle motion along the optical axis. Scale bar, 1 *µ*m. (b) Simultaneous, interlaced iNTA–fluorescence characterization of a mixed nanoparticle sample containing 30 nm gold nanoparticles, 40 nm red FluoSpheres, and 100 nm green FluoSpheres. (Left) iNTA scatter plot reveals distinct clusters corresponding to the three particle populations, color-coded by their fluorescence signatures. Diamond-shaped symbols indicated two data points from unidentified impurtiy particles. (Right) Per-particle fluorescence intensity scatter plot (488 nm vs. 561 nm). Dot symbols represent matched iNTA–fluorescence measurements, whereas grey crosses indicate values obtained from temporally uncorrelated iNTA and fluorescence frames, serving as a control for random co-localization (see Methods). The dashed lines mark the transition to negative intensities in the green and red channels, caused by median background subtraction.

### Fluorescence quantification of immunolabelled extracellular vesicles (EVs)

EVs are small, membrane-enclosed particles that are released into the extracellular space and are expected to play key roles in intercellular communication.^27^ Their molecular cargo reflects the physiological state of the cell of origin, making EVs promising candidates for non-invasive diagnostics. For example, tumor-derived EVs carry distinct oncogenic RNA and protein signatures, enabling early cancer detection.^28–31^ iNTA measurements have proven very helpful in characterizing the EV size distribution and in providing valuable information about their effective refractive index. This has allowed one to distinguish and categorize different particle populations.^32,33^ Augmenting this ability with fluorescence information provides access to biochemical specificity, which is crucial for medical research. For instance, variations in surface protein composition can reflect differences in biogenetic origin and functional specialization. In particular, tetraspanins CD9 and CD81 have been recognized as informative surface markers for probing EV subpopulations.^19^

To demonstrate the performance of our platform for this application, we labelled HEK293 EVs with anti-CD9–Alexa Fluor 488, anti-CD81–DyLight 550, or both antibodies. For quantification of the number of dye molecules carried by individual EVs, we first characterized surface-immobilized samples. Here, we examined the photophysical properties of the antibody-conjugated dyes using stepwise bleaching of surface-immobilized antibodies and antibody-labeled EVs. This calibration enabled the conversion of fluorescence intensities into the number of dye molecules per particle. Based on these estimates, EVs were classified as fluorescence-positive using a threshold of one estimated dye molecule, corresponding to a false-positive rate of approximately 1%.

In Figure 4, we present our results for singly and doubly-labeled EVs. In each case, we first show the conventional iNTA plot to characterize the size distribution (Figure 4a,d,g) and then present the fluorescence signals on a logarithmic scale (Figure 4b,e,h). We find that in the case of singly-labeled EVs, 32% were CD9-positive (Figure 4b) and 48% were CD81-positive (Figure 4e), consistent with the ensemble-averaged western blot analysis. In Figure 4h, we show that dual-labeling experiments further resolve how these markers co-occur on individual EVs. The largest shared population consisted of EVs that were positive for both CD9 and CD81 (28%), while a substantial fraction carried CD81 in the absence of detectable CD9 (22%). In contrast, EVs that were only positive for CD9 constituted a minor subpopulation (4%), indicating the asymmetric distribution of these tetraspanins across the EV population. Furthermore, we found a size-dependent increase in the fluorescence intensity, implying that larger EVs possess a higher abundance of tetraspanin.

**Figure 4:**
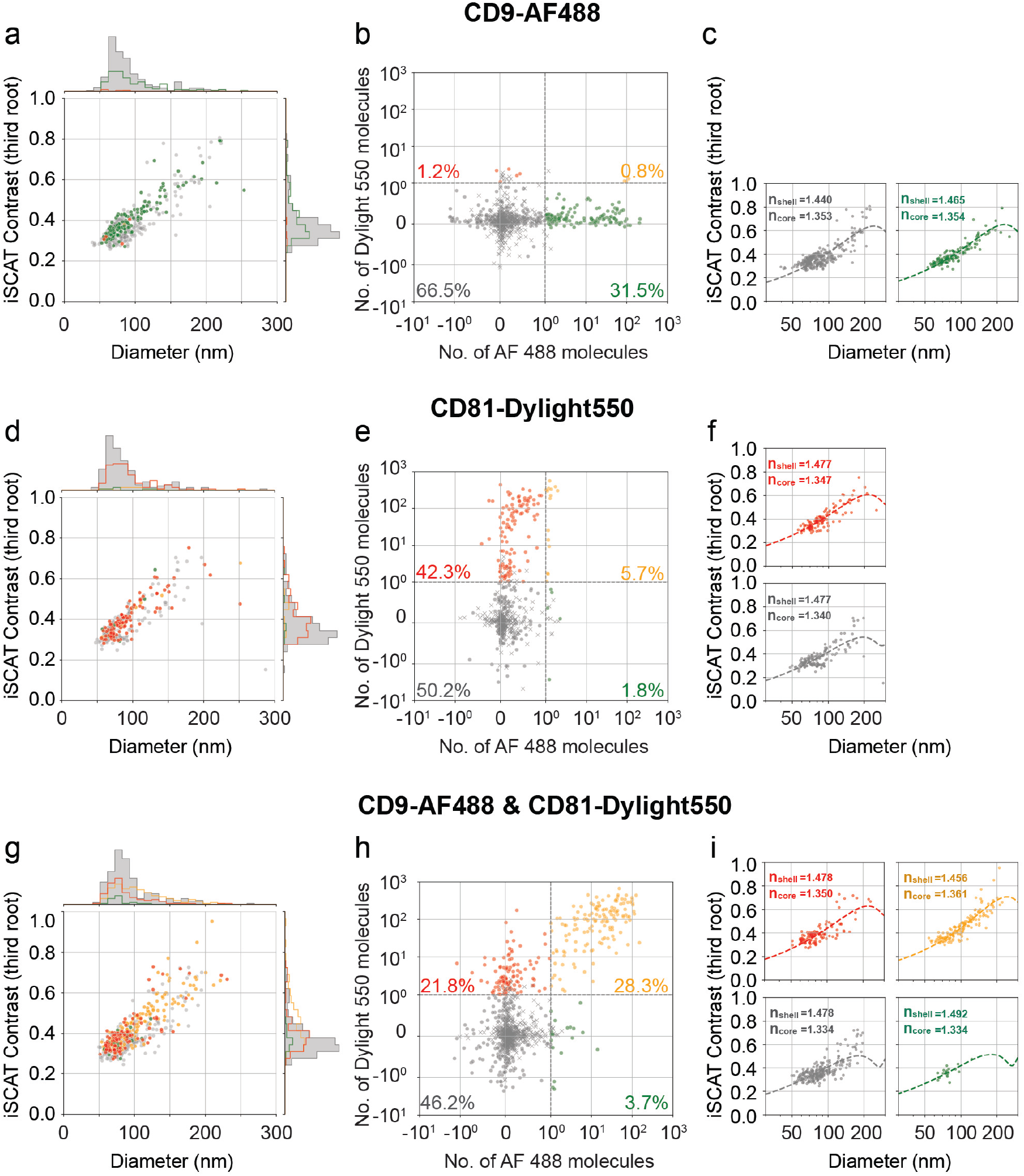
Molecular profiling of HEK293 EVs by iNTA-F. (a, d, g) Conventional iNTA scatter plots of the third root of the iSCAT contrast and diameter measured from diffusion trajectories. Fluorescent EVs are color coded. Particles without fluorescence signal are shown in grey. (b, e, h) Two-color fluorescence quadrant plots showing estimated AF 488 and DyLight 550 molecules for EVs labelled with CD9–AF488, CD81–DyLight 550, and both antibodies. (c, f, i) Channel-separated iNTA scatter plots for each labelling condition. Fluorescence-defined subpopulations are shown in separate quadrants. Grey: fluorescencenegative; Red: DyLight 550; Green: Alexa Fluor 488; Orange: double-positive.

In Figure 4c,f,i, we map the fluorescence information onto the EV size and iSCAT contrast at the single-particle level. Across all labeling conditions represented by symbols of different colors, the overall iNTA distributions of labeled EVs closely resembled those of unlabeled control samples. This shows that antibody labeling did not globally alter the detectable scattering properties of EVs.

Next, we present conventional iNTA plots in Figure 4c,f for fluorescence-positive populations (green or red symbols) as well as the fluorescence-negative sub-population (grey symbols). In addition, the orange symbols in Figure 4i represent the population that was doubly labeled. Interestingly, we observe a general trend that the spread in each fluorescence-positive subpopulation follows more closely the distribution expected for core-shell vesicles, shown by the theoretical solid curve. The observed trend might suggest that some EVs have different constitutions in terms of their biochemical inner load as well as their protein and sugar coating,^34^ which might in turn affect the efficiency of various labeling strategies. The platform introduced in this work is ideally suited for in-depth investigation of such important nuances.

## Conclusion

In this work, we extended iNTA measurements by integrating multicolor fluorescence detection with single-molecule sensitivity. This enables simultaneous measurement of particle size, refractive index, concentration, and quantification of fluorophore copies on freely diffusing particles. While, this information might also be obtained in other approaches,^35–38^ we believe iNTA-F presents a particular advantage in combining sensitivity and high throughput in a single platform. Although we chose to test the performance of our methodology on EVs, the approach is applicable to a broad range of nanoparticle systems, including synthetic nanoparticles,^39^ lipid-based drug carriers,^40^ viral particles,^41^ and other heterogeneous assemblies where size, composition, and surface functionalization play an important role. ^42^ The ability to correlate physical and molecular properties at the single-particle level also paves the way for biomarker-based sorting strategies. ^43^ Increasing the number of excitation wavelengths and incorporating sequential label-exchange strategies would increase multiplexing capacity and enable the analysis of more complex biomarker panels.^44^

## Methods

### Measurement Setup

We used three laser at wavelengths 638 nm, 488 nm and 561 nm for illumination in iNTA and excitation of fluorescence in the green and red channels, respectively. Laser beams were co-aligned and directed to the sample. The iSCAT signal was recorded on a CMOS camera, capable of detecting small variations on a large background signal, while fluorescence emission was spectrally filtered, split into two channels, and sent to separate sCMOS cameras for measuring weak signals. The frame rate for iSCAT imaging was set to 5000 fps, while fluorescence imaging ran at 100 (fluorosphere detection) and 10 (liposomes and EVs) frames per second. The iSCAT and fluorescence cameras were temporally synchronized with each other and the fluorescence lasers using a custom-built Arduino circuit.

To minimize photobleaching during fluorescence measurements, a flow system was implemented to continuously replenish fluorescent samples within the field of view. A six-channel glass-bottom flow chamber (ibidi, µ-Slide VI 0.5 Glass Bottom, 80607) was used, with each channel measuring 17 mm in length, 0.54 mm in height, and 3.8 mm in width. Fluid flow was controlled by a syringe pump (LA-30), ensuring stable and reproducible sample exchange throughout the measurement period. To reduce nonspecific binding of biological particles within the flow chamber, the channel surfaces were passivated using silane-PEG.

### Trajectory analysis and fluorescence quantification

Recorded videos were processed using median background correction and radial variance transform as previously described.^45^ Particle tracking was performed with ‘trackpy’ Python package,^46^ retaining only trajectories with more than 500 localizations. Particle size was determined from the diffusion constant.^14^ The effective refractive index (RI) was subsequently calculated using the extracted particle size and measured the iSCAT contrast.^14^ Fluorescence videos were background-corrected using median background over 5 s before and after the recording. They were spatially aligned to iSCAT videos using fluorospheres deposited on a clean coverglass. To account for the illumination inhomogeneity, fluorescence images were divided by the normalized laser illumination profile.

The trajectories extracted from iSCAT videos were split into segments with length given by the ratio of the frame rates fps_iSCAT_*/*fps_fluo_, so that each portion corresponded to a single frame of the fluorescence camera. Each segment was used to generate a boolean mask. This was done by drawing a circle with radius 1px at the position of each localization and adding all the circles together. In order to extract the total fluorescence counts per particle, the fluorescence frame was multiplied by the boolean mask, and the counts were added together. The maximum value of pixel counts was extracted for each trajectory.

As a control for background fluorescence and potential false-positive signal assignments, we implemented frame-mismatched analysis between iNTA trajectories and fluorescence image sequences. In this procedure, particle trajectories extracted from the iNTA channel were correlated with fluorescence frames acquired at non-corresponding time points, thereby eliminating any true temporal association between particle positions and fluorescence emission. In samples with elevated background fluorescence, such as those containing freely diffusing unbound dye molecules, this mismatched analysis yields spurious fluorescence intensities when trajectories are incorrectly correlated with fluorescence fluctuations. The resulting intensity distributions, therefore, provide a direct estimate of background-driven false-positive signals. Only samples for which fluorescence intensities obtained from frame-mismatched analysis were consistent with the noise floor were considered for quantitative fluorescence assignment. Fluorescence signals derived from time-matched analysis were attributed to particle-associated emission rather than background contributions.

### Single-molecule dye preparation and imaging

1 mg of Alexa Fluor 488 NHS Ester (A20000, Thermo Fisher Scientific) or Dylight550 NHS Ester (62262, Thermo Fisher Scientific) was dissolved in 500 µL of dimethylformamide (DMF) and subsequently diluted 10^8^-fold in LC-MC Grade H_2_O (Merck, 115333100). Then, 30 µL of this solution was spin-coated onto a pre-cleaned coverslip (Zeiss, 474030-9010-000). The spin coating speed was set to 3000 rpm for 30 s. The coverslip was pre-cleaned by sonication in 2% Hellmanex III (Hellma, 9-307-011-4-507) for 5 min, followed by rinsing with Milli-Q H_2_O. It was further sonicated in ethanol and then in isopropanol for 5 min each. Next, the coverslip was blow dried by nitrogen. The autofluorescence of the coverslip was checked prior to spin coating with dye molecules.

### Formation of liposomes and lipid staining

Liposomes were prepared following previously established methods to closely resemble the characteristics of EVs. The lipid composition (mol %) of the liposomes used in our experiments was DSPC/Chol/DOPS (40/40/20 mol %). Briefly, the desired amounts of lipids (Avanti) were mixed in chloroform, and the mixture was dried under a stream of nitrogen gas to create a thin, homogeneous lipid film. This film was further dried overnight under vacuum desiccation to remove any residual solvent. The dried film was then hydrated with Milli-Q H_2_O at 70°C. The hydrated lipid solution was sonicated for 10 min and then passed 37 times through an extruder equipped with a 100 nm polycarbonate filter to form liposomes.

For liposome staining with lipid dyes, the size, concentration, and refractive index of the liposomes were first measured using iNTA. Pre-diluted liposomes (1 × 10^10^ particles/mL) were stained with MemBright 488 (Merck, SCT083; final dilution 1:1,000) by incubating with the dyes at 37 °C for 1 hour on an orbital shaker at 300 rpm. Excess dye was removed using a PD SpinTrap G-25 spin column (Cytiva, 28918004) following the manufacturer’s protocol prior to measurement on iNTA.

### HEK293 EV production and isolation

HEK293 cells were cultured in DMEM culture medium supplemented with 10% Fetal Bovine Serum (FBS) in 4-6 T175 flasks until they reached 80% confluency. The medium was then replaced with FBS-free DMEM, and the cells were cultured for an additional 48 hours. Following this, the cell culture medium was collected and centrifuged at 300g for 5 min to remove cells and debris. The supernatant was further centrifuged at 10,000g (Beckman Coulter, Optima XPN-80 Ultracentrifuge) for 35 min at 4°C to pellet larger particles. The resulting supernatant was then subjected to ultracentrifugation (Beckman Coulter, Optima XPN-80 Ultracentrifuge) at 150,000g for 2 hours and 5 min at 4°C to pellet smaller extracellular vesicle (sEV). This pellet was washed with phosphate-buffered saline (PBS) and spun again at 150,000g for 2 hours and 5 min at 4°C. Finally, the pellet was resuspended in 100 µL of PBS, aliquoted, and stored in a -80°C freezer until further use.

### Antibody labeling with dye and detection of antibody-labelled EVs

Anti-CD81-Dylight550 and anti-CD9-Alexa488 were used to label HEK293-EV. Briefly, 10 µL of EV (1 × 10^10^ particles/mL, diluted in PBS and quantified by iNTA) were incubated with 4 µL of anti-CD81-PE and/or 4 µL of anti-CD9-Alexa488. The total volume was adjusted to 20 µL with PBS. The mixture was incubated at 37 °C for 1 hour with continuous orbital shaking at 300 rpm. Excess antibodies were removed using a qEVsingle size-exclusion chromatography column (IZON, ICS-35). Fractions 3 to 6 (totaling 680 µL) were collected, diluted as needed, and loaded into the syringe pump for measurement.

For detection of antibody-labeled EVs, both the 488 nm and 561 nm fluorescence channels were recorded at 10 Hz exposure time, while the iNTA channel was acquired at 5000 Hz. For interlaced excitation, the 488 nm and 561 nm lasers were alternately switched on at 5 Hz each, whereas the 638 nm laser for the iNTA channel remained continuously on. Each sample was measured for 30 min.

## Acknowledgement

The authors thank Simone Ilhoff and Fabian Greiner for their technical assistance. We are also grateful to Patricia Bassereau, Stephan Götzinger, Tim Hebenstreit, Siwei Luo and Hisham Mazal for valuable discussions and insights.

